# Symbolic regression for empirically realistic population dynamic time series

**DOI:** 10.64898/2026.02.16.706224

**Authors:** Cheyenne N. Jarman, Taal Levi, Mark Novak

**Author notes:** **Code Availability** All code is available at https://github.com/cheyennejarman/SymbReg_KelpForest_Simulations. **Author contributions** CNJ and MN project ideation; CNJ data generation and analysis; CNJ, TL, and MN writing.

## Abstract

Applications of machine learning in ecology are rapidly expanding. Symbolic regression is gaining particular attention for its success in reverse-engineering human-readable explanatory population models, including the logistic growth and Lotka-Volterra equations, from simulated and laboratory-based population time series. However, field-based populations often lack the characteristics of the idealized time series used in prior assessments. We evaluated the utility of symbolic regression for such time series by quantifying its success for synthetic data varying in sampling density, population cycle asymmetry, process noise, and the erroneous consideration of spurious variables. We further compared two data preprocessing options for estimating population growth rates, and four evaluation workflows for selecting equations. Results indicate that a trade-off between sampling density and process noise primarily drives equation and variable recovery. Symbolic regression failed to recover the underlying equation at sampling densities below 10 points per cycle; however, at higher sampling densities, process noise did increase equation recovery rates. Importantly, although the true model was frequently recovered at sampling densities of 25 or more points per cycle, it was not consistently selected by the evaluation workflows. This discrepancy highlights a need for more robust post-algorithm selection criteria to identify the focal equation among competing candidates.

## Introduction

Ecologists are motivated by the goal of describing patterns and process through observation, experimentation, and mathematical modeling. Mathematical models are particularly important for abstracting high-dimensional systems into equations that bring focus to key variables and mechanisms. For over a century, such models have been constructed based on *a priori* conceptualizations. For example, the variables, parameters, and formulations of the logistic growth (Verhulst, 1838) and Lotka-Volterra (Lotka, 1925; Volterra, 1926) models, as well as the myriad other population dynamics models descended from these, were specified by logic, intuition, and first principles (Hilborn & Mangel, 1997). Unfortunately, although this approach may work for well-studied or relatively simple systems, it is susceptible to biases stemming from preconceived notions, paradigms, and scientific inertia (Bonnaffé *et al*., 2021; Ellner *et al*., 2002; Kuhn, 1970; Novak *et al*., 2025). In recent years, several approaches have therefore begun to take a more data-driven approach to dynamical model building (Solomatine & Ostfeld, 2008). Tools like neural networks (Bonnaffé *et al*., 2021), empirical dynamical modeling (Munch *et al*., 2020; Sugihara & May, 1990), and symbolic regression (Bongard & Lipson, 2007) in particular are increasingly being applied to time series to infer the data-generating processes from the data, rather than specifying a particular model or set of models *a priori*.

As a form of scientific machine learning (Baker *et al*., 2019), symbolic regression stands out among these data-driven approaches for its goal of providing human-readable, mechanistically-interpretable equations in the spirit of the classical equations of theoretical ecology (Gerwin, 1974; Langley, 1977). Given data on a response variable and a set of putative predictor variables, symbolic regression algorithms use a process akin to biological evolution to evolve generations of equations through mutation, the recombination of parental equation components, and the selection and transmission of equations based on their fit to the data. (See the *Supplementary Materials* for an overview.) The first ecological application of symbolic regression to population time series appears to have been Bongard & Lipson (2007) who found it to successfully recover the two-species Lotka-Volterra competition equations even when the simulated time series were subject to observation noise. They also found it to produce similar equations and mechanistic interpretations to previous *a priori* models of Lynx-Hare interactions when applied to the classic Hudson Bay population time series. Gaucel *et al*. (2014) extended these findings to demonstrate symbolic regression”s ability to recover the Lotka-Volterra predator-prey equations from simulated time series subject to observation noise under sampling strategies in which the time elapsed between observations was either regular or irregular. This was followed by Martin *et al*. (2018) who demonstrated symbolic regression”s success in reverse-engineering the logistic growth equation, the Lotka-Volterra predator-prey equations, and a stage-structured population model from empirical time series of single and paired predator-prey laboratory populations. Finally, Chen *et al*. (2019) assessed the method”s ability to recover and reverse-engineer the population growth and nonlinear functional response equations of predator-prey populations from simulated and laboratory time series, focusing specifically on the data”s dynamical informativeness through a contrast of time series exhibiting a stable limit cycle versus a transient approach to a fixed-point equilibrium. Although symbolic regression was only successful at recovering the generative equation in the cycling case, Chen *et al*. (2019) concluded that its failure in the transient case was a limitation of the data rather than of symbolic regression itself. Altogether, the current literature therefore provides a unanimous endorsement of symbolic regression for learning population dynamic models from time series data.

Despite prior demonstrations of its success, symbolic regression”s ability to reverse-engineer the equations of more complex ecological time series with field-relevant sampling characteristics remains unknown. While symbolic regression offers a powerful framework for recovering mechanistic models, it has yet to be fully operationalized for generating insights from empirical systems. This lag in broad-scale application persists because we lack a clear understanding of how the method handles the reality of field data compared to idealized simulations. In this regard it is particularly unclear whether the sampling densities of typical population time series are sufficient for symbolic regression to succeed. Prior assessments have assumed sampling intervals many times shorter than typical population cycle period lengths or generation times (e.g., Inchausti & Halley, 2001). Further knowledge gaps remain regarding time-series preprocessing practices (i.e., the use of time series interpolation methods) and how process noise rather than observation noise, the symmetry of population cycles, and the erroneous consideration of spurious predictor variables (whereas previous benchmarks were restricted to *a priori* known drivers) affects the method”s reliability. Finally, there has been little consideration of the alternative methods by which the suite of equations returned by symbolic regression can be assessed to identify and select the correct or “best” equation.

By generating empirically relevant time series from a mechanistic model of giant kelp populations, we evaluated symbolic regression”s success in regards to each of these factors in six case studies, with each subjected to five different sampling densities. We included a contrast of four alternative evaluation workflows for selecting candidate equations from along the Pareto frontier, the set of equations representing the optimal trade-off between complexity and explanatory power. We also examined the diversity of returned equations along and behind the frontier to gain insight into the effectiveness of each workflow. We structure our manuscript by introducing our generative population model, further motivating and describing the case studies, and detailing our implementation of symbolic regression and means of quantifying its rates of success for each workflow. Overall, we identified significant limitations under certain field-relevant conditions, but determined the subset of conditions under which performance is acceptable, which can serve as a guide for ecologists to determine when symbolic regression is suitable for their inferential problems.

## Methods

### The generative model

We used the Bence & Nisbet (1989) delay-differential model for giant kelp (*Macrocystis pyrifera*) to generate population time series. This model qualitatively reproduces the cycles observed in empirical giant kelp populations by encapsulating an effect of adult kelp on the recruitment of juvenile kelp in competing for space. The model describes the population growth rate of adult kelp *A* at time *t* as

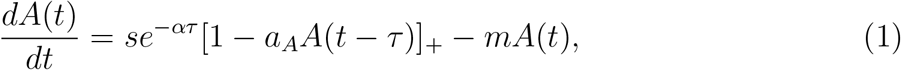

where *s* represents the juvenile settlement rate, *α* and *m* represent the juvenile and adult per capita mortality rates, *a*_*A*_ represents the proportion of space occupied by an adult individual, and *τ* represents the temporal lag of juveniles maturing to adults (i.e., juveniles are represented implicitly by a time-delayed effect of adult population size). The operator [*x*]_+_ = *x* if *x* ≥ 0 else 0 provides an abrupt stop in growth once no space is available. We modified this model to include the possibility of multiplicative process noise in juvenile settlement and adult mortality (i.e., *s* → *s* exp[*ϵ*_1_] and *m* → *m* exp[*ϵ*_2_] with *ϵ*_*i*_ ~ 𝒩 (0, *σ*_*i*_)). Previous studies have only considered unbiased observation noise (i.e., measurement error), the effect of which diminishes with increased sampling density and time-series replication, while neglecting process noise (i.e., the inherent stochasticity of population dynamic processes). Notably, although process noise may increase the challenge of reverse engineering equations just like observation noise, it may also make time series more informative of their underlying processes by pushing systems to explore a larger space of variable states and conditions (Boettiger, 2018).

We chose the Bence-Nisbet model for several additional reasons as well. First, because it can generate both symmetrical and asymmetrical population cycles in a biologically justified manner. Many real-world populations exhibit asymmetric cycles (e.g., a rapid increase followed by a slow decay (Turchin, 2003)), yet with the exception of Chen *et al*. (2019), all prior studies have considered only symmetrically cycling populations. This may be important because the combination of asymmetry and low sampling density can bias estimated rates of population change due to the higher chance of capturing the longer, slower phase, particularly under sampling regimes that are regular (i.e., periodic), as most field-based time series are. Second, because it is more complex than previously used generative models by virtue of a time-delayed variable and use of exponential and []_+_ operators. Its inclusion of a time-delayed variable was of particular interest because (*i*) many population time series entail time-delayed processes arising from higher-dimensional multi-species and age- or stage-structured processes (De Roos & Persson, 2003; Turchin, 2003), (*ii*) several time-series methods (e.g., empirical dynamical modeling) rely on the representation of multi-dimensional systems using time-delay embedding (i.e., Takens’ (1981) theorem), and (*iii*) it provided a means to introduce non-trivial spurious variables in our assessments; all prior studies provided their symbolic regression algorithms only with variables known to influence dynamics, which is an unlikely scenario in field contexts.

### Case studies

#### Time series generation

We used the model to generate time series for six case studies, evaluating both a discrete-time and continuous-time approach under symmetrically and asymmetrically cycling deterministic time series (cases 1–4; Fig. 1a-d) and a continuous-time approach on asymmetrically cycling stochastic time series under low and high levels of process noise (cases 5–6; Fig. 1d-f). The juvenile maturation rate was set to *τ* = 2 following Bence & Nisbet (1989) for all case studies. Most other parameter values were specific to each case study (Tables S.1-S.2). Each time series was simulated for 1000 time steps, from which the last 15 complete population cycles were retained. The cycle period lengths were 5.5 and 13.5 time steps in the symmetric and asymmetric deterministic cases, and averaged 13.2 and 12.5 time steps for the symmetric stochastic cases under low and high process noise, respectively. To generate the actual time series that we provided to symbolic regression, we downsampled each case study”s time series at sampling densities of 100, 50, 25, 10, or 5 time points per cycle. Although not reported, prior studies used synthetic time series with sampling densities of hundreds of time points per cycle and laboratory time series with sampling densities no lower than approximately 12 time points per cycle.

**Figure 1:**
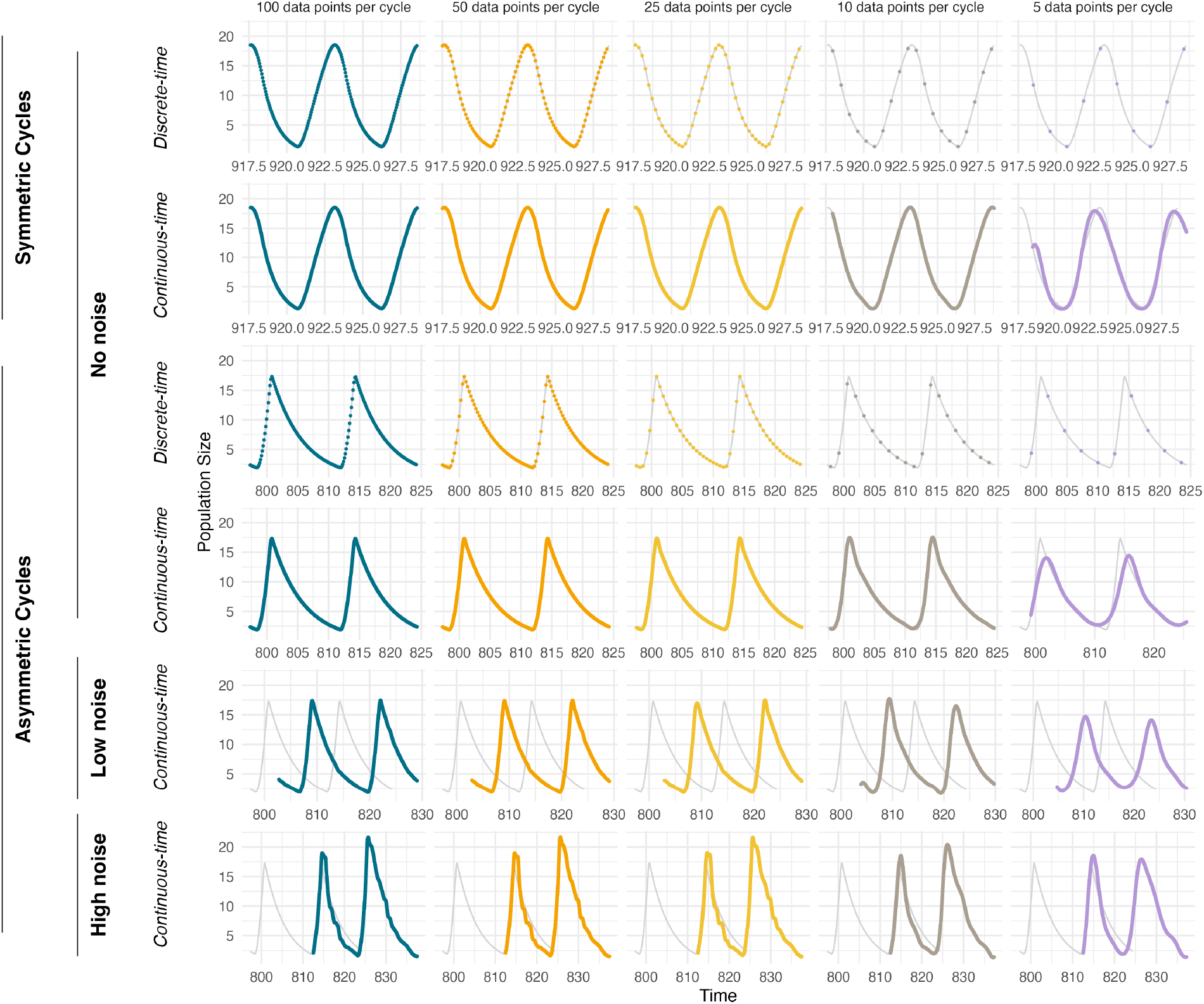
Time series illustrating our six case studies and the consequences of varying sampling density within each. A total of 15 cycles were used in our analyses, but only two are plotted for visualization. The light gray lines show the dynamics of the instantaneous deterministic Bence-Nisbet model for reference. The overlaid colored points show the (downsampled and interpolated) data from which the per capita growth rates and (lagged) putative predictor variables were obtained and provided to symbolic regression.

#### Response and predictor variables

Each downsampled time series was used to generate the response and predictor variables that we provided to symbolic regression. As the response variable, we used the per capita growth rate, estimated differently depending on the use of a discrete- or continuous-time approach. For the discrete-time approach we calculated the per capita growth rate using the log-ratio of successive abundances, ln (*A*_*t*+Δ*t*_*/A*_*t*_) */*Δ*t*, as this converges on the instantaneous per capita growth rate 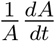 as Δ*t* decreases (see *Supplementary Materials*). For the continuous-time approach we calculated the first derivative of the cubic spline interpolated log-transformed abundances (using SciPy, Virtanen *et al*., 2020) since 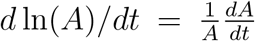. Our comparison of these two approaches was motivated by prior simulation- and laboratory-based studies that have used continuous-time preprocessing, relying on sufficiently high sampling densities to avoid issues associated with alternative interpolation techniques. Our use of log-transformed population size differs from prior studies as it avoided the interpolation of negative population sizes to which the use of natural-scale population sizes is susceptible when sampling density is low.

Previous studies provided their algorithms with state variables that were known to pertain directly to the focal dynamic. Similarly, we conducted a preliminary study to ensure that the symbolic regression algorithm we used could recover the generative equation when only *A*(*t*) and *A*(*t* − 2) were considered as putative predictors (see the *Supplementary Materials*). However, in empirical contexts the true causal variables are typically not known. Our analyses therefore also considered the extraneous predictor variables *A*(*t* − 1) and *A*(*t* − 3). Note that only *A*(*t*) and *A*(*t* − 2) appear in the generating equation (eqn. 1), making *A*(*t* − 1) and *A*(*t* − 3) autocorrelated but spurious determinants of the dynamics. For both the discrete-time and continuous-time approaches, the lagged variables were determined by back-transforming interpolated log-transformed population sizes.

### Symbolic regression

#### Implementation

Prior ecological studies used custom-built symbolic regression algorithms. We used PySR (v 1.3.1; Cranmer, 2023), a Python library with a Julia backend that has seen frequent use in Physics. Unlike prior algorithms, PySR evolves equations using multiple semi-independent populations of equations between which migration can occur in order to explore a greater volume of equation space (see the *Supplementary Materials* for further details). With mean squared error (MSE, a.k.a. L2-loss) as the metric of equation fitness, we used 96 populations and ran each downsampled time series through PySR 100 independent times. Each of the 6 × 5 × 100 = 3000 searches was run on a single compute node of a high performance computing cluster equipped with 64 CPU cores and a total of 514 GB of RAM for 22 hours or until an equation reached an MSE of 2 · 10^−9^, though in almost all cases the search time was exhausted first. Running 100 independent searches allowed us to quantify rates of equation recovery in a probabilistic manner rather than the binary success/failure manner of prior studies.

#### Equation and variable recovery success

We compared four workflows (two subjective and two objective) to select candidate equations from among the learned equation and assess symbolic regression”s ability to recover the Bence-Nisbet model. Success rates were quantified in two ways for each workflow: as the proportion of 100 runs in which the selected equation (*i*) included only the two correct predictor variables of the Bence-Nisbet model (i.e., *A*(*t*) and *A*(*t* − 2)), and (*ii*) included only the two correct predictor variables and matched the Bence-Nisbet model in its functional form and parameter values (to three significant digits).

Prior studies assessed success using several approaches, including by subjectively inspecting the Pareto frontier of the learned equations to search for the true or a recognizable equation. The Pareto frontier represents the equations that maximize fitness (e.g., minimize MSE) for their given level of complexity; it reflects the most efficient trade-off between equation fitness and complexity. PySR quantifies an equation”s complexity as the total count of all parameters, variables, and operators it contains (which for the Bence-Nisbet model is 14). Typically, the simplest model that provides the greatest increase in fitness along the Pareto frontier is selected as the best performing model, although a user”s goals may cause the selection of other models. For Workflow 1, we followed this practice by visually inspecting plots of MSE versus complexity and selecting the simplest equation that affected the largest additive decrease in MSE relative to all other equations along the frontier. Workflow 2 repeated this visual inspection using ln(MSE) to select the simplest model affecting the largest multiplicative decrease in MSE relative to all other equations along the frontier. PySR itself provides an objective, quantitative selection criterion equivalent to Workflow 2 called *score* that is calculated as the negative discrete log loss change per unit change in complexity (Cranmer, 2023), which is what we used for Workflow 3. In Workflow 4, we used the Bayesian Information Criteria (BIC) for model selection (Schwarz, 1978). BIC evaluates equation performance by penalizing the equation”s fit (its MSE) by the count of its parameters and the sample size of the data (Hilborn & Mangel, 1997; Höge *et al*., 2018). Next to AIC, BIC remains the most commonly used information criterion but has the property of being asymptotically consistent as the sample size increases (i.e., will select the true model — rather than the most efficient model — when it is present). For all workflows, we only considered equations with a complexity of up to 20.

#### Explanatory variable diversity and selection success

In addition to quantifying rates of equation and variable recovery success for each workflow as above, we also (*i*) looked for the Bence-Nisbet model and (*ii*) characterized the diversity of variable combinations present within the top (lowest MSE) equations of each complexity level returned by PySR, combining all 100 runs of each case-study sampling-density combination. We did the latter by counting the number of unique variable combinations that occurred in the up to the top 10 equations, restricting ourselves to equations in the complexity range of 10–18. (While we considered up to 10 equations, fewer were recovered for certain complexities across the 100 searches. Because these instances were rare, we maintain the “top 10” terminology throughout for brevity.) Without necessitating an equation selection workflow, this analysis was motivated by the context in which a user does not know the true driving variables or is not expecting classical population models to apply — hence would not look for or be able to identify these along the Pareto frontier or in any suite of returned equations — but could gain confidence and insight by virtue of symbolic regression returning a low diversity of equations that combined just a few dominant variables at or near the Pareto frontier.

Finally, in our most judicious assessment aligned with those of several prior studies, we determined whether the Bence-Nisbet model appeared in any of the returned equations of a given complexity in the range of 10–20, combining all 100 runs of each case study sampling-density combination. This permitted us to distinguish the ability of symbolic regression to evolve the correct generating equation from the ability of the four workflows to identify it along the Pareto frontier. Thus, when the Bence-Nisbet model did occur, we determined its maximum MSE-rank by counting the number of equations of the same complexity that exhibited a lower MSE than it.

## Results

### Equation and variable recovery success

We observed widely varying success rates (0–75% of 100 runs) in recovering the Bence-Nisbet model using the four equation selection workflows (Fig. 2). Overall, success rates were most influenced by sampling density and the presence of process noise, with no substantive or consistent effects of cycle symmetry versus asymmetry or preprocessing method. Although none of the workflows proved reliable, the two objective workflows (3 and 4) generally performed more poorly than the two subjective workflows (1 and 2) when success was observed. The highest rates of success were observed when applying Workflow 1 to the continuous-time preprocessed asymmetric time series having high process noise, subject to sampling densities of 50 or more time points per cycle. Modest rates of success (10–20%) were also observed when applying Workflows 1 and 2 to otherwise equivalent time series having low process noise, subject to sampling densities of 50 or more time points per cycle. However, rates of success were very low (*<* 10%) or zero for the vast majority of other case studies, workflows, and sampling densities. In fact, the Bence-Nisbet was never recovered by any workflow when sampling densities were less than 50 time points per cycle, except for a few instances in the presence of process noise.

**Figure 2:**
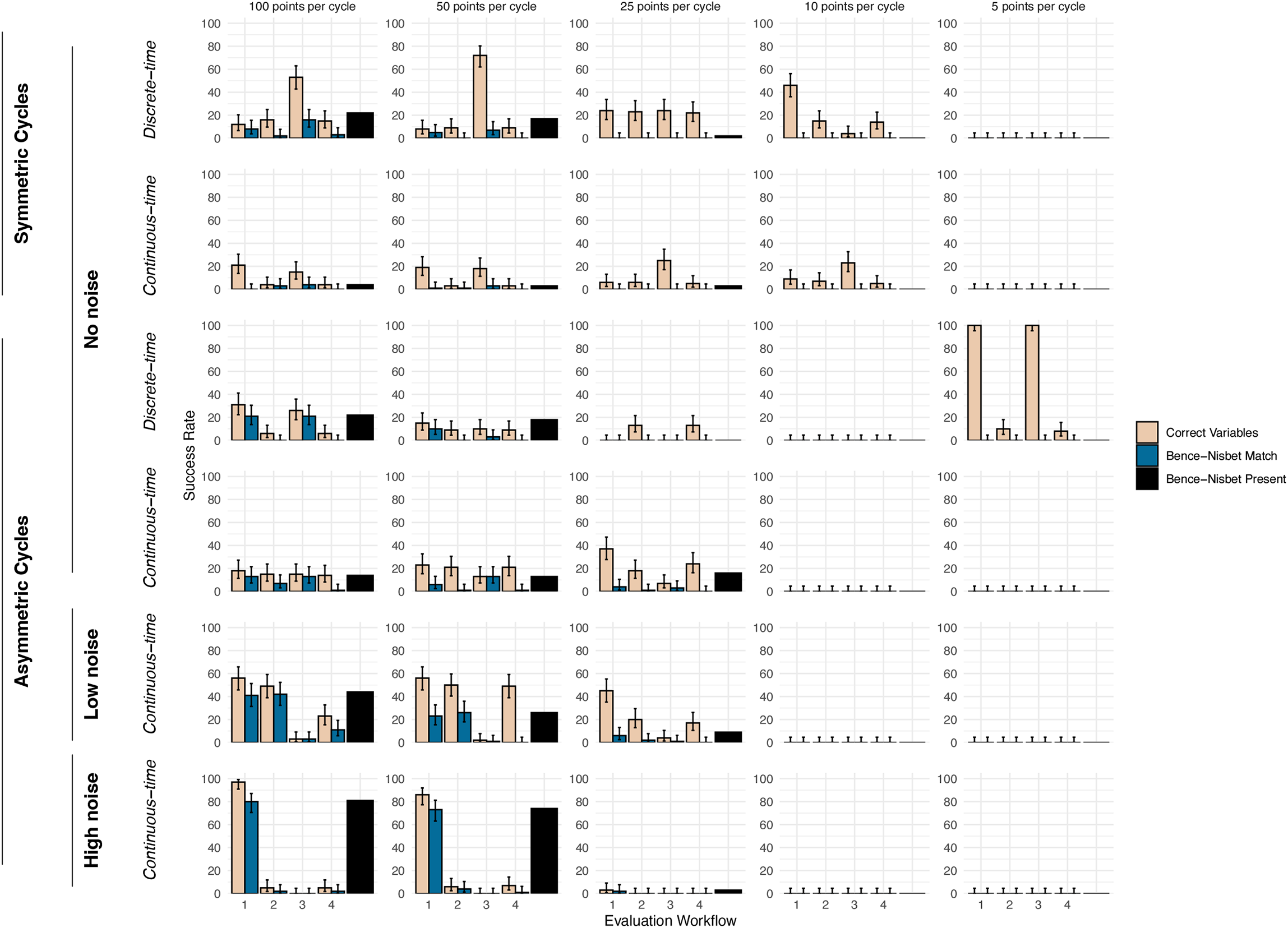
The success rate of symbolic regression and the four model selection workflows in selecting equations that contained the correct variables or that contained the correct variables and matched the formulation of the Bence-Nisbet model with which we generated the data. Confidence intervals reflect binomial uncertainties given a sample size of 100 independent runs. The black bars indicate the proportion of the 100 independent runs in which the Bence-Nisbet model occurred, regardless of whether a workflow selected it or whether it occurred in the top 10 equations.

We observed higher — albeit still low and widely varying (0–100% of 100 runs) — rates of success in using the workflows to select equations which contained only the correct variables of the Bence-Nisbet model along the Pareto frontier (Fig. 2). A rate of 100% correct variable recovery occurred anomalously for Workflows 1 and 3 in the case of discrete-time preprocessed deterministic asymmetric cycles subject to a sampling density of only 5 time points per cycle. (None of these equations matched the Bence-Nisbet model”s structure, see below.) Similarly, an anomalously high rate of correct variable recovery occurred when applying Workflow 3 in the case of discrete-time preprocessed deterministic symmetric cycles at a sampling density of 50 time points per cycle. Beyond these exceptions, rates of correct variable recovery generally paralleled the patterns observed in the equation recovery success, tending to be low overall (*<* 20%) and higher in the presence of process noise when applying Workflow 1. Unlike for equation recovery, correct variable recovery success rates remained above zero at sampling densities less than 50 time points per cycle in several deterministic cases.

### Explanatory variable diversity and selection success

The diversity of variable combinations among the top 10 (lowest MSE) equations at each complexity level was low for a given case study (Fig. 3). No set of the top 10 equations contained more than 7 of the 11 possible pairwise-or-higher combinations of variables that could occur at complexities above 9, with agreement among all top equations for a single combination of variables or just two combinations of variables being by far the most frequent (48.7% and 27.7% of 267, respectively). Out of the 2581 equations examined in this analysis, *A*(*t*) and *A*(*t* − 2) occurred most frequently (31.8%), followed by *A*(*t*), *A*(*t* − 1) and *A*(*t* − 2) (23.2%). For most case studies, the frequency of equations that contained only the two correct variables, *A*(*t*) and *A*(*t* − 2), increased substantially at all complexity levels as sampling density increased, with the majority of the top equations per complexity containing only the two correct variables in several high sampling density case studies. (The notable exceptions were the case study considering deterministic symmetric cycles preprocessed with the continuous-time approach, for which the frequency of the correct variable equation remained low for all sampling densities, and the previously mentioned anomalous case of discrete-time preprocessed deterministic asymmetric cycles at a sampling density of 5 time points per cycle.) For all case studies, sampling densities less than 50 points per cycle tended to result in equations containing spurious variables, with the frequency of equations containing the combinations of *A*(*t*), *A*(*t* − 1) and either *A*(*t* − 2) or *A*(*t* − 3) increasing particularly in the presence of process noise. Equations containing *A*(*t*), *A*(*t* − 1), and *A*(*t* − 2) specifically also occurred more frequently for the two symmetric cycle case studies at high sampling densities. Equations that included only the two spurious variables, *A*(*t* − 1) and *A*(*t* − 3), were rare, except in the two cases of deterministic asymmetric cycles (discrete- and continuous-time preprocessed) at a sampling density of 10 points per cycle.

**Figure 3:**
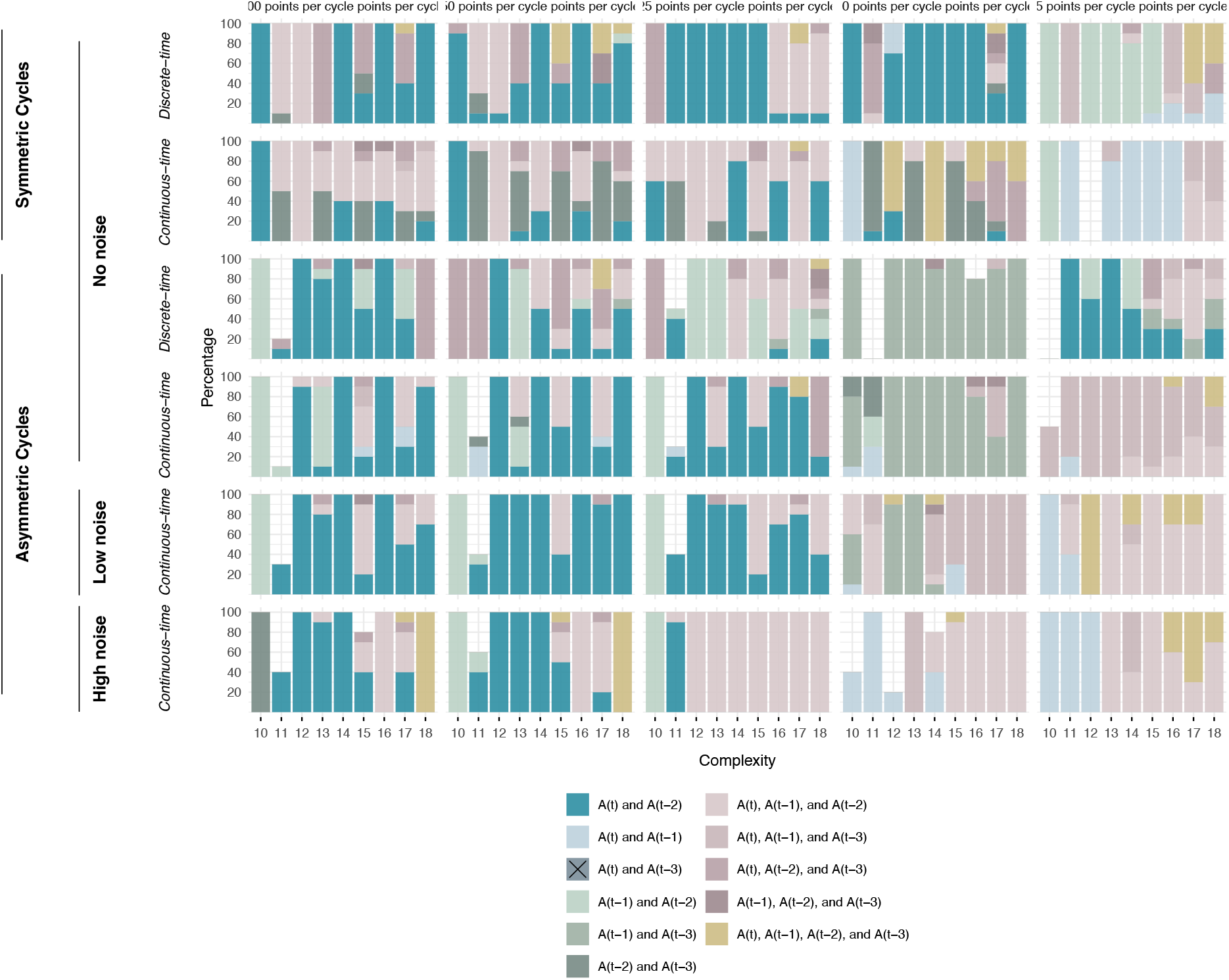
The proportional frequency of variable combinations present among the up to top 10 (lowest MSE) equations of each complexity level in the range of 10–18 (combining all 100 runs of each case-study sampling-density combination). The Bence-Nisbet model with which the data were generated has a complexity of 14 and contains only *A*(*t*) and *A*(*t* − 2), making the inclusion of *A*(*t* − 1) and *A*(*t* − 3) spurious. All possible variable combinations are depicted in the legend; however, the combination of *A*(*t*) and *A*(*t* − 3) never occurred.

Despite not being consistently selected from along the Pareto frontier of any given run using the four workflows, the Bence-Nisbet model was evolved by symbolic regression for nearly all case studies (black bars in Fig. 2), and occurred in the top 10 (lowest MSE) equations of all case studies, given a sampling density of 25 or more points per cycle (Fig. 4). In the case studies in which the Bence-Nisbet model occurred within the top 10 as complexity 14, it had the lowest MSE, ranking first among all returned equations, 60% of the time. However, in four instances it held a lower rank (higher MSE), and in two cases it failed to appear entirely, not placing along the Pareto frontier (Fig. 5). In most of the cases where the Bence-Nisbet model occurred, equations of higher complexity which were equivalent to the Bence-Nisbet model (i.e., equations that had extraneous constants that could be removed by algebraic simplification) also occurred (Fig. 4). These equations often also ranked first at their complexity level, but were more likely to drop in rank and out of the top 10 as their complexity increased (Fig. 5).

**Figure 4:**
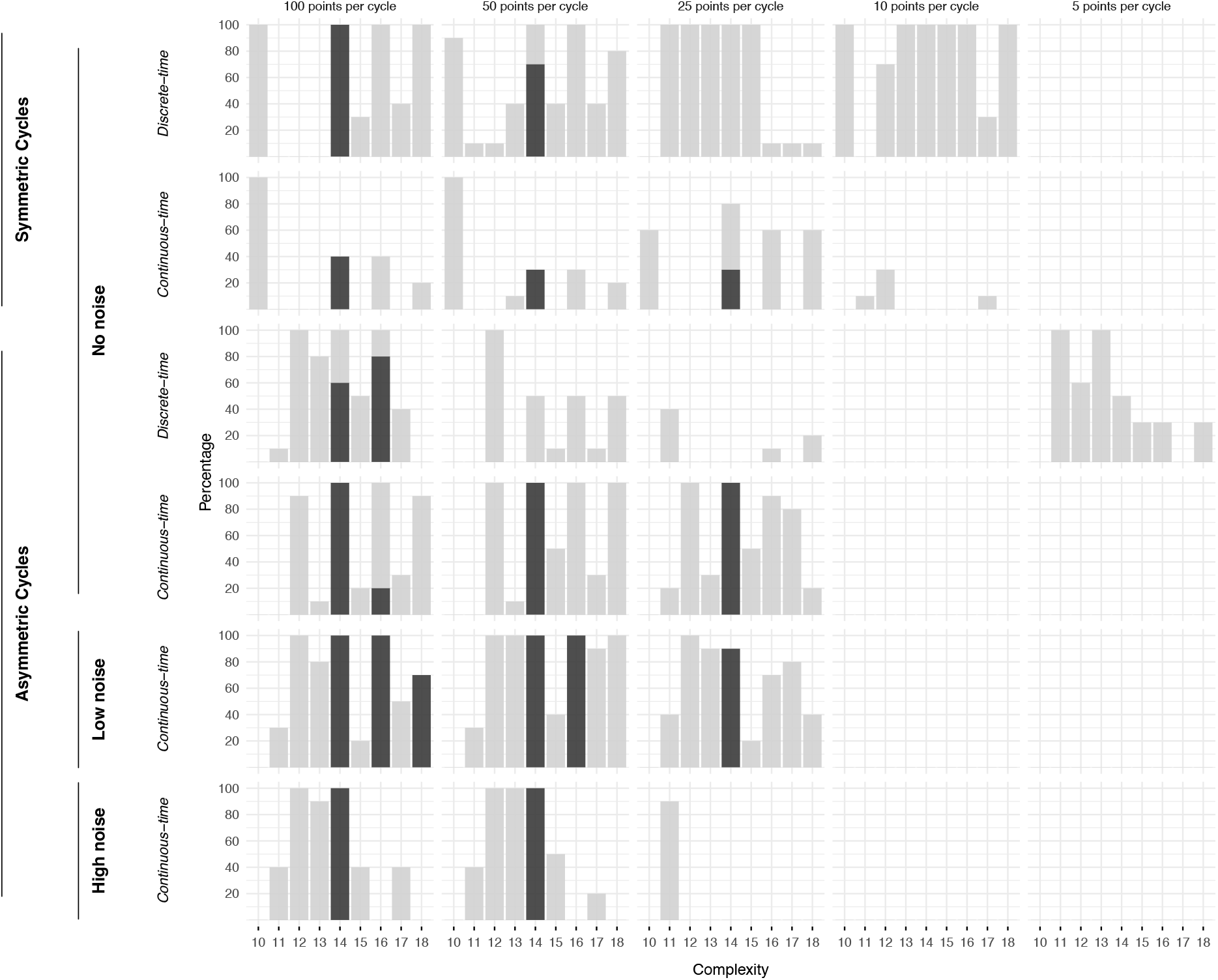
The proportion of the top 10 (lowest MSE) equations that correctly matched the Bence-Nisbet model in each complexity level in the range 10–18 (combining all 100 runs of each case-study sampling-density combination). Lightgray backgrounds indicate instances in which at least one equation occurred that contained only the two variables of the Bence-Nisbet model, *A*(*t*) and *A*(*t* − 2), as shown in Fig. 3. The complexity level of the Bence-Nisbet model is 14. Matches at complexities greater than 14 occurred when the equations could be algebraically simplified to the Bence-Nisbet model. Note that matches at odd-valued complexity levels are not possible given the even-valued complexity of the Bence-Nisbet model and the necessity of including an operator for every additional parameter or variable.

**Figure 5:**
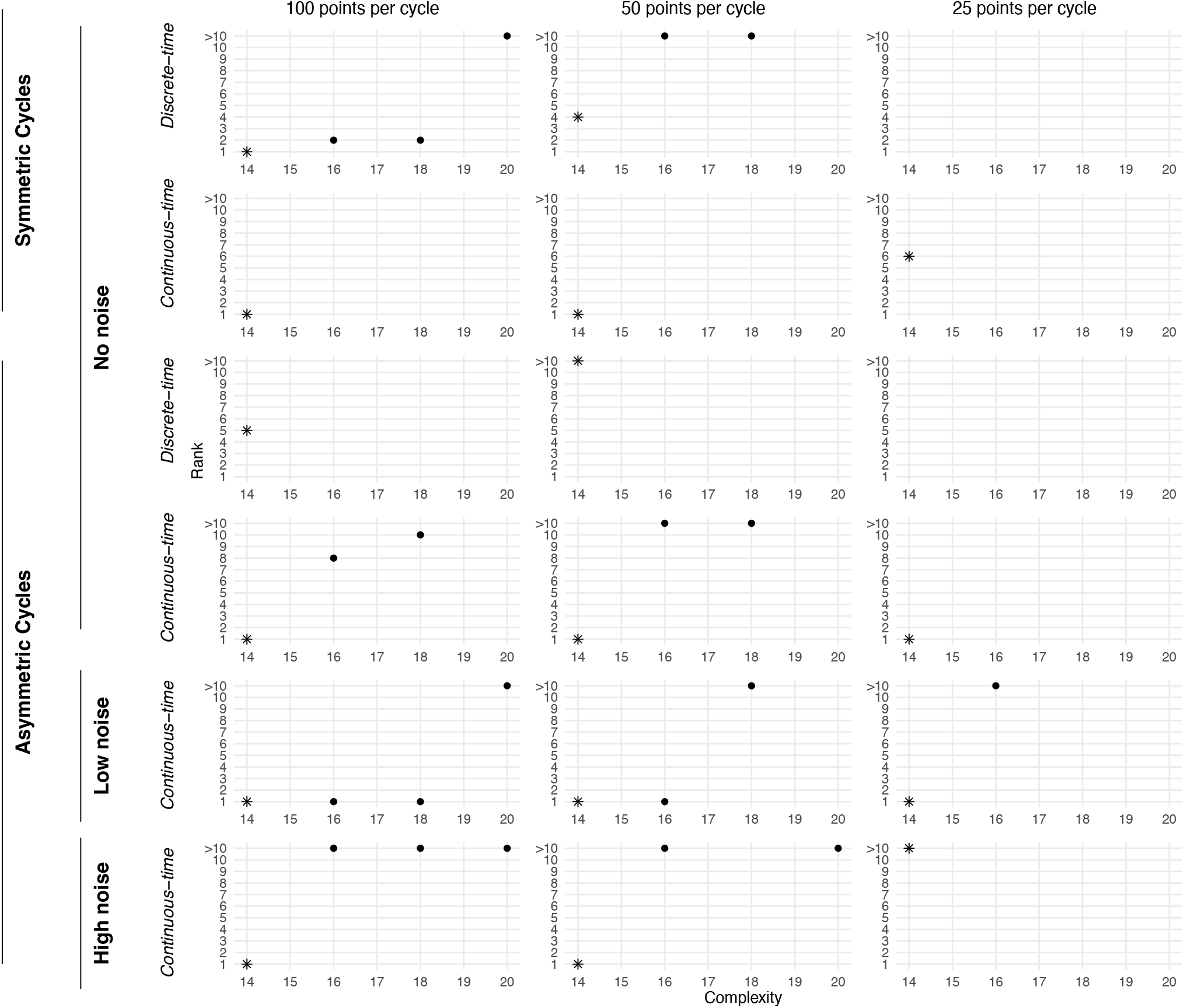
The maximum MSE-rank of the Bence-Nisbet model when it was recovered, reflecting the number of equations that exhibited a lower MSE than it in the set of up to 100 equations returned by the 100 runs of each case-study sampling-density combination. The complexity level of the Bence-Nisbet model is 14 (indicated by asterisks), hence lower levels are not possible. Ranks at complexities greater than 14 (indicated by points) occurred when the equations could be algebraically simplified to the Bence-Nisbet model. The Bence-Nisbet model was never recovered for time series of 10 and 5 points per cycle or for the deterministic discrete-time processed case study at 25 points per cycle (see Fig. 4).

## Discussion

We evaluated the utility of symbolic regression for recovering the generative population model of synthetic time series that varied in sampling density, process noise, cycle asymmetry, and preprocessing approach, and assessed four workflows for selecting candidate equations from among those returned. In contrast to prior studies based on densely sampled time series, our results indicate that both equation recovery and equation selection are substantially less reliable under characteristics typical of field-based time series. In particular, recovery success depended strongly on sampling density and, to a lesser extent, on the presence of process noise, while population cycle asymmetry and preprocessing approach had comparatively little influence. Even when the true equation was evolved, none of the four workflows proved consistently reliable at identifying it.

Nevertheless, our findings also suggest reasons for cautious optimism. Indeed, a central insight of this study is the need to distinguish among three related but distinct components of symbolic regression analyses: the algorithm”s ability to evolve the correct equation, the position of that equation relative to and along the Pareto frontier, and the performance of workflows in selecting an equation. Although the generative model we used was more complex than those of prior studies, symbolic regression frequently did evolve the correct equation at least once across repeated runs when sampling density was sufficiently high. In these cases, the correct equation often occurred among the top-performing equations of its complexity level and typically ranked first or near-first in terms of MSE across all runs. Moreover, the diversity of equations near the Pareto frontier was generally low and dominated by models containing the correct variables. Together, these results underscore the value of long-term ecological time series of high density and indicate that symbolic regression can recover meaningful mechanistic structure from them, but also reveal that identifying this structure requires improved criteria for robust equation selection.

### Equation generation versus equation selection

The low rate of successful recovery on a per-run basis raises the question of whether failures arose from limitations of our symbolic regression implementation or from limitations of the data and evaluation workflows. Our results suggest that the latter two, rather than computational constraints, were the primary factors. All searches were run for substantially more runs, longer durations, and across more populations of equations than in prior ecological studies, making it unlikely that recovery failures reflect insufficient exploration of equation space.

Instead, symbolic regression frequently evolved incorrect equations that fit the data as well as or better than the true model. This outcome is consistent with extensive prior work demonstrating that structurally distinct models can generate indistinguishable dynamics when data do not sufficiently constrain the mechanism (e.g., Novak & Stouffer, 2021; Perretti *et al*., 2013; Song & Levine, 2025). In such cases, equation selection criteria that rely on fit and various measures of complexity alone cannot be depended upon to identify the true equation, even when it is present. In addition, in our study, the correct equation not only occurred behind the Pareto frontier but more commonly was present on the frontier, yet was not selected by an equation selection workflow, particularly for the workflows that were metric-based and hence entirely objective. Together, these findings highlight the need for criteria that better reflect structural identifiability rather than goodness-of-fit and traditional measures of complexity alone (e.g., Novak & Stouffer, 2021). Approaches that explicitly consider equations both on and behind the Pareto frontier, or that incorporate additional dynamical diagnostics like the ability to reproduce key statistical measures of the time series (e.g., Song & Levine, 2025), may also help to improve inference.

### Sampling density, process noise, and data informativeness

Sampling density exerted the strongest influence on symbolic regression”s performance. At sampling densities below 25–50 points per cycle, the correct equation was rarely evolved or reliably selected. Because all our time series were sampled with densities determined on a per-cycle basis and the lowest assessed sampling frequency exceeded the Nyquist-Shannon frequency of the simulated dynamics, differences in recovery across case studies are unlikely to reflect classical aliasing effects (see the *Supplementary Materials*). Instead, reduced sampling density appears to limit recovery by diminishing data informativeness, affecting the accuracy of interpolated abundances and the estimation of growth rates, and consequently the ability of the data to discriminate among alternative equation variables and structures.

Our analyses therefore emphasize that sampling density must be considered relative to a system”s dynamical frequencies or characteristic timescales when applying equation-discovery methods. For strongly seasonal systems, sampling once or even twice a year, as is often the case for field-based population time series, will not be enough. Because achieving high sampling density is often challenging in field-based contexts, alternative strategies — such as irregular sampling (Gaucel *et al*., 2014), compressive sampling (Candes & Wakin, 2008), adaptive sampling (Henrys *et al*., 2024), or the use of state-space rather than rate-versus-state equation formulations that avoid the error-sensitive estimation of growth rates (Munch *et al*., 2020) — may be useful in mitigating these constraints and warrant further investigation.

A more favorable result for field-based contexts was that process noise consistently increased recovery success. Indeed, as increasingly recognized in other contexts (Boettiger, 2018; Coulson *et al*., 2004; Dennis *et al*., 2006), stochastic perturbations appear to enhance data informativeness by expanding coverage of the system”s attractor (see the *Supplementary Materials*) and revealing responses to variation that are absent with deterministic cycles. These effects likely reduce the set of dynamically equivalent models and help distinguish among competing equations. That said, contrary to our expectations, the highest noise level that we assessed often performed as well as or better than the low-noise case. Identifying the threshold at which process noise transitions from informative to detrimental thus remains an important direction for future work.

### Cycle asymmetry, preprocessing, and variable identification

Population cycle asymmetry had little effect on symbolic regression”s performance, with asymmetric cycles performing comparably to or slightly better than symmetric ones. This result contrasts with our expectation that asymmetric cycles would be more vulnerable to under-sampling, but suggests that asymmetry alone does not strongly interact with sampling density to constrain recovery when sampling density is sufficiently high on a per-cycle basis.

Similarly, we observed no consistent advantage of either discrete-time or continuous-time preprocessing for estimating per capita growth rates. Although both approaches can introduce distinct sources of error, these effects were generally secondary to those of sampling density and process noise. Stronger differences may emerge under alternative dynamics or more extreme asymmetries.

Introducing spurious predictor variables revealed that symbolic regression can correctly identify the true driving variables when sampling density is sufficiently high, even when the true equation structure is not evolved or identified. At lower sampling densities, however, auto-correlation among lagged predictors likely favored the inclusion of spurious but dynamically redundant variables. This pattern suggests that variable selection failures at low sampling densities again arise from data informativeness rather than from an inherent limitation of symbolic regression. The implementation of methods like convergent cross-mapping to remove spurious variables during pre-processing may be useful for improving inference (Munch *et al*., 2020; Sugihara & May, 1990). That said, the observation that the combination of *A*(*t*), *A*(*t* − 1), and *A*(*t* − 2) specifically was the second most common combination of variables is particularly interesting given that it reflects the variables by which the true dynamics of the our presumed model may be faithfully represented by time-delay embedding using Takens” (1981) theorem. Future work should examine the degree to which selection biases arise when spurious state variables differ in their degree of autocorrelation with the system”s true driving variables, as well as the extent to which time-delay embedding may interfere with the identification of a system”s true causal structure. Additionally, investigating the frequency of specific equation structures, rather than just variable occurrences, would be valuable; however, such an effort will necessitate automated structural analysis given the high dimensionality of the search space.

## Conclusions

Our results indicate that symbolic regression can recover mechanistically meaningful population models, but only under conditions of suffciently informative data and with careful attention to equation selection. Given suffcient sampling, the primary limitation identified here lies not in equation generation but in identifying the correct model from among dynamically equivalent alternatives. Improving post hoc evaluation criteria — particularly those that move beyond exclusive reliance on the Pareto frontier — thus represents a key opportunity for methodological advancement.

## Acknowledgments

We thank Bruce Menge, Dave Lytle, Rebecca Hutchinson, the Model Enabled Machine Learning group, and affiliates of the Novak lab for thoughtful discussion and feedback.

## Supplementary Materials for

### A Symbolic regression and PySR

#### Symbolic regression

Symbolic regression aims to find simple yet accurate equations to explain a given dataset. Using a process akin to biological evolution, generations of equations are evolved, where individual equations with a better fit to the data have a higher chance of producing offspring equations in the following generation. Symbolic regression begins with the creation of an initial generation of equations. Two equations with high fitness are selected as “parental equations”. These parental equations produce an offspring equation for the next generation of equations through additions, deletions, recombinations, or mutations of their components. Multiple pairs of parental equations are selected within any given generation. Additionally, equations from previous generations may move forward into subsequent generations. This equation evolution process occurs for a set amount of time or generations. This algorithm can be implemented in various ways – the four previous ecological studies utilized custom-built algorithms. We implemented symbolic regression with PySR (Cranmer, 2023), an open-source Python library with a Julia backend. We will describe symbolic regression through the PySR lens, noting that while there are overarching similarities in the workflow, specific implementation details differ from other algorithms.

PySR begins with the user providing a list of predictor variables, a single response variable, and a suite of mathematical operators (both built-in and custom, user-defined operators) to combine terms (Fig. S.1a). PySR differs from other symbolic regression algorithms in that evolution can occur within and between populations of equations (Fig. S.1b). The number of populations is pre-specified as a user-input argument. PySR initiates the specified number of populations, each containing an initial generation of equations created through random assemblages of predictor variables, parameters, and mathematical operators. Within each population, equation evolution occurs through tournament selection, where the population is randomly subsampled and the individuals with the highest fitness values are selected as parental candidates (Cranmer, 2023). Mean squared error (MSE) is commonly used as the fitness metric, thus the equation with the highest fitness will have the lowest MSE. The two parental candidates exchange subexpressions of their equation to produce a new equation via crossover. Mutation can also occur and only needs a single parental candidate. In this case, a subexpression of the parental equation is modified (e.g., a change in a parameter value or the operator used). Equation evolution is also done between populations (i.e., gene flow) where equations from one population randomly migrate to another population and vice versa, allowing for potentially new equation genes to enter a given population (Fig. S.1b) (Cranmer, 2023). PySR”s output is a file containing the best-performing equations and the respective MSE across a range of complexity levels, where complexity is defined as the sum of the numbers of parameters, variables, and operators present in a given equation (Fig. S.1c). To help with equation selection, the user can plot the equation”s MSE against the complexity level (Fig. S.1d).

**Figure S.1:**
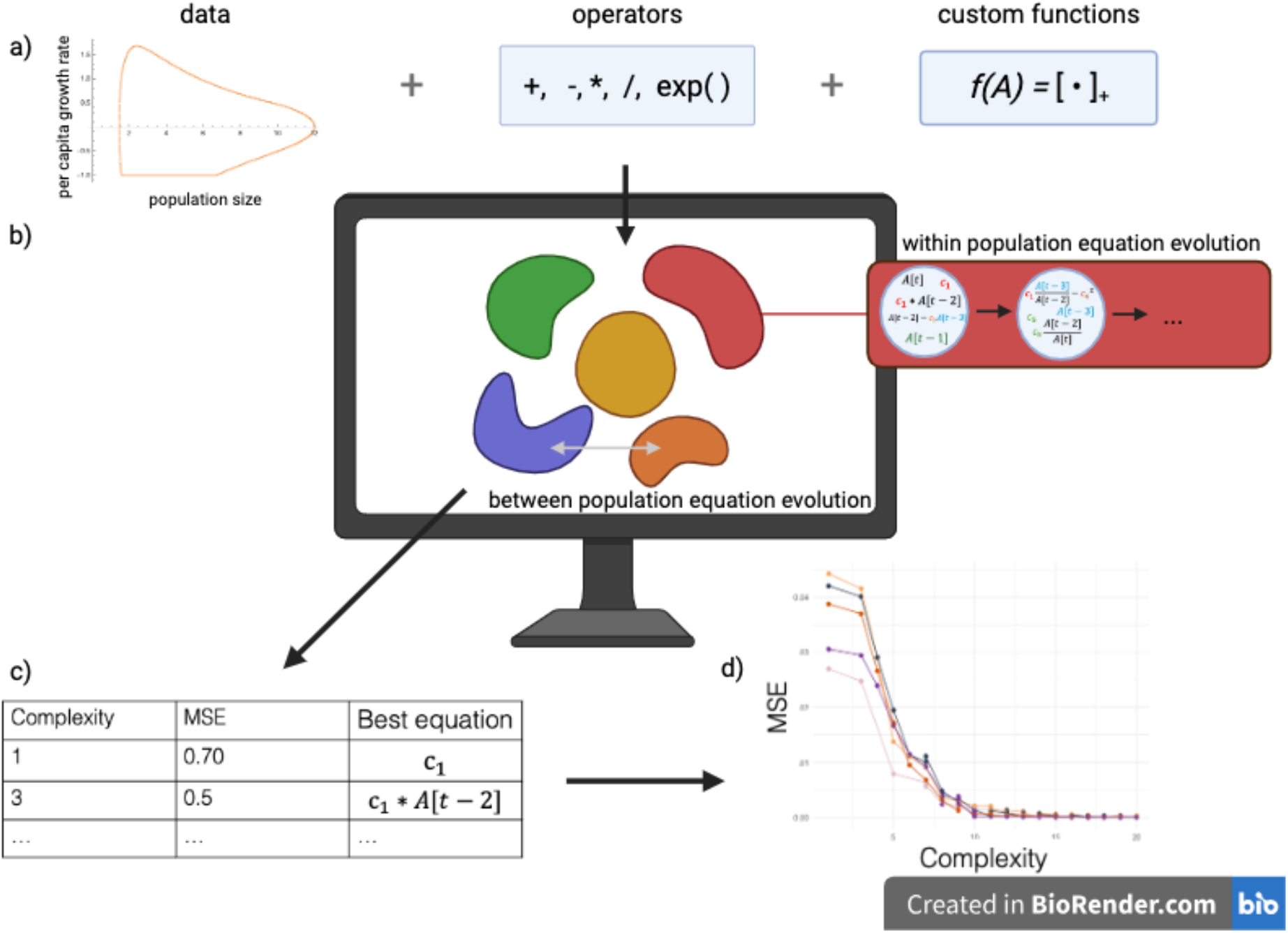
An overview of the symbolic regression workflow. (a) The inputs provided to the symbolic regression include the data, mathematical operators, and any custom functions. (b) Equation evolution occurs at two stages, (*i*)“within population evolution” wherein multiple equations exist within a single generation and parental equations are selected based on fitness level to create a new equation in subsequent generations and (*ii*)“between population evolution” wherein new equation components can enter a population by migration between populations. (c) The output at the end of a run entails a list of the best (lowest MSE) equation for each level of complexity, along with its MSE values and complexity level. (d) These results may be visualized to discern the Pareto frontier of equations by plotting the complexity level on the x-axis and MSE on the y-axis.

#### PySR

PySR allows for the specification and customization of many different arguments. We specified that the following operators could be used: addition, subtraction, multiplication, division, and the exponential function. We also created a custom function that matched the non-linear function, []_+_. This was the only custom function we used. PySR”s *populations* argument defines how many populations of equations are evolving. More populations will increase the diversity of equations and run time (Cranmer, 2023). We used 96 populations in all analyses. The user can also specify how many iterations, or the total number of generations, PySR will run for. The PySR documentation recommends setting this argument to a high value, so we specified a maximum of 1 · 10^10^ iterations. Finally, the user can control how much time PySR spends searching by specifying the timeout argument. We specified our search to run for 22 hours. Equation overfitting and bloating are known concerns for symbolic regression algorithms. PySR manages these concerns by defining a maximum equation size (user-specified or default value 30, we used the latter). Additionally, PySR ensures relatively even equation complexity representation by tracking the frequency of particular complexity levels and the complexity levels of the most recent equations (Cranmer, 2023). While there is no clear recommendation for the optimal number of runs needed to ensure the global optimum is reached, we ran each simulated data set through PySR independently 100 times. Previous studies also conducted (but combined) multiple search runs, with the number ranging from 3–30.

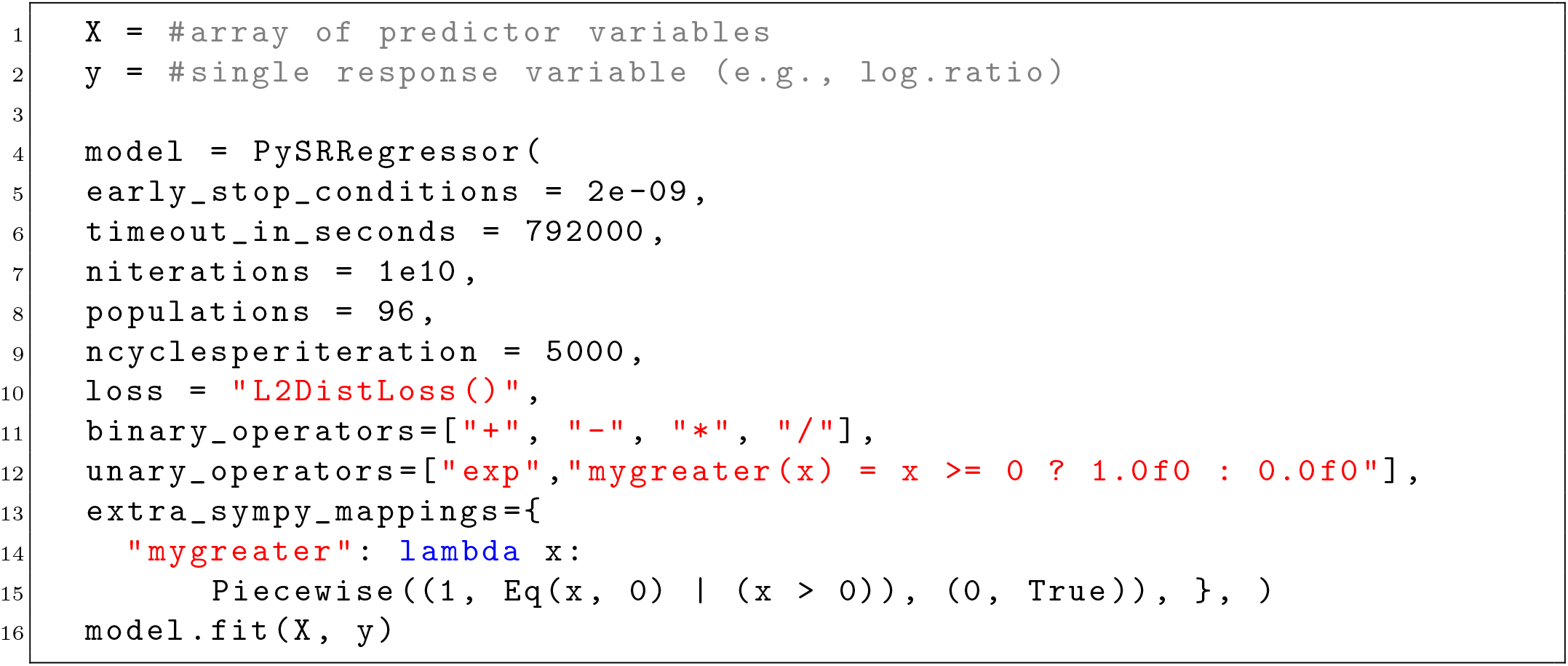

### B Parameters used to simulate time series

Our modifiication of the Bence & Nisbet (1989) model incorporated process noise as

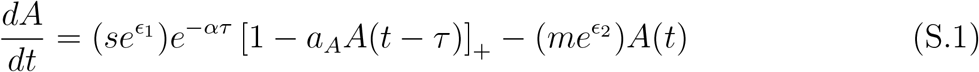

where *ϵ*_1_ and *ϵ*_2_ are random variables with *ϵ*_*i*_ ~ 𝒩 (0, *σ*_*i*_) and all parameter values are given in Tables S.1 & S.2.

**Table S.1:** Parameter values used to generate time series.

| Parameter | Symbol | Symmetric | Asymmetric |
| --- | --- | --- | --- |
| Juvenile settlement rate | $s$ | 55 | 55 |
| Juvenile mortality rate | $\alpha$ | 0.35 | 0.15 |
| Space occupied by individual adult | $a_A$ | 0.1 | 0.35 |
| Adult mortality rate | $m$ | 1 | 0.2 |
| Juvenile maturation delay | $\tau$ | 2 | 2 |

**Table S.2:**
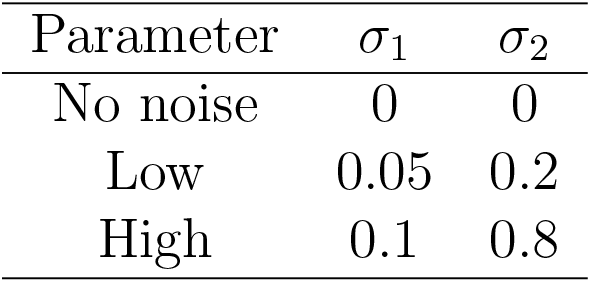
Parameter values used to introduce process noise.

| Parameter | $\sigma_1$ | $\sigma_2$ |
| --- | --- | --- |
| No noise | 0 | 0 |
| Low | 0.05 | 0.2 |
| High | 0.1 | 0.8 |

### C Time series period lengths and aliasing

The period length of cycles produced by the model for each of the four cases was determined by Fourier analysis using *SciPy*. In the deterministic cases, the period lengths were 5.5 and 13.5 for the symmetric and asymmetric cases, respectively. For the stochastic cases, we simulated the time series for 1000 time steps and took the last 500 time steps to average the period length, which was 13.2 and 12.5 time steps for the symmetric stochastic cases under low and high process noise, respectively.

Since in every case we examined the cycles had periods of at least five sampling intervals — i.e., we observed a minimum of five time points per cycle — their fundamental frequencies were well below the Nyquist-Shannon cutoff (Shannon, 1949). Consequently, the down-sampled series are unlikely to suffer aliasing; only if very high-frequency harmonics exceeded half the sampling rate would they have risked being reflected at lower apparent frequencies.

### D Asymptotic equivalence of discrete and continuous per capita growth rates

The log ratio 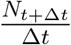 is asymptotically equivalent to the instantaneous per capita growth rate 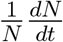 as Δ*t* approaches 0. That is,

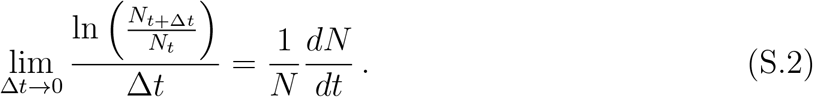

This may be shown as follows.

Using the quotient rule for logs, we can rewrite the log ratio as the difference between *N*_*t*+Δ*t*_ and *N*_*t*_,

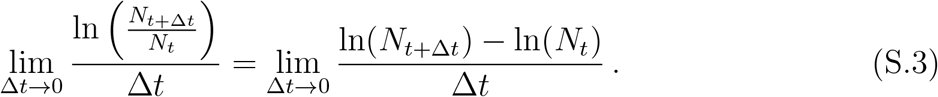

The term on the right-hand side is equivalent to the definition of a derivative, thus we can rewrite the expression as

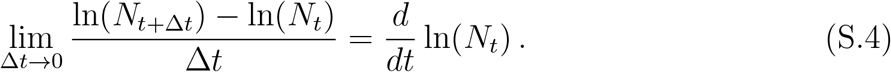

Lastly, applying the chain rule shows how the time derivative of the logarithm of population size relates to the instantaneous per capita growth rate. Specifically, since ln(*N*_*t*_) is a function of *N*_*t*_, which itself varies with time, we differentiate ln(*N*_*t*_) with respect to *N*_*t*_ and then multiply by the rate at which *N*_*t*_ is changing over time. Because the derivative of ln(*N*_*t*_) is 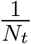, this yields

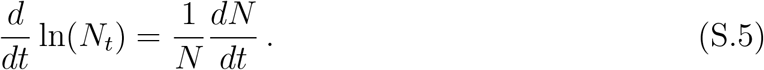

### E Equation recovery when using only *A*(*t*) and *A*(*t*−2)

We simulated the Bence-Nisbet model using parameter values that generated asymmetric cycles (as in Table S.1) and provided PySR with data downsampled to a sampling density of 100 time points per cycle with the per capita growth rate using either the continuous-time or the discrete-time preprocessing approach. We performed 100 independent runs of PySR for each time series. For each run, we evaluated all resulting equations and ranked them according to their MSE. We then examined the rankings to determine whether the Bence-Nisbet model was present (Fig. S.2). In both the continuous- and discrete-time approaches, the Bence-Nisbet model was recovered among the top 10 ranked models.

**Figure S.2:**
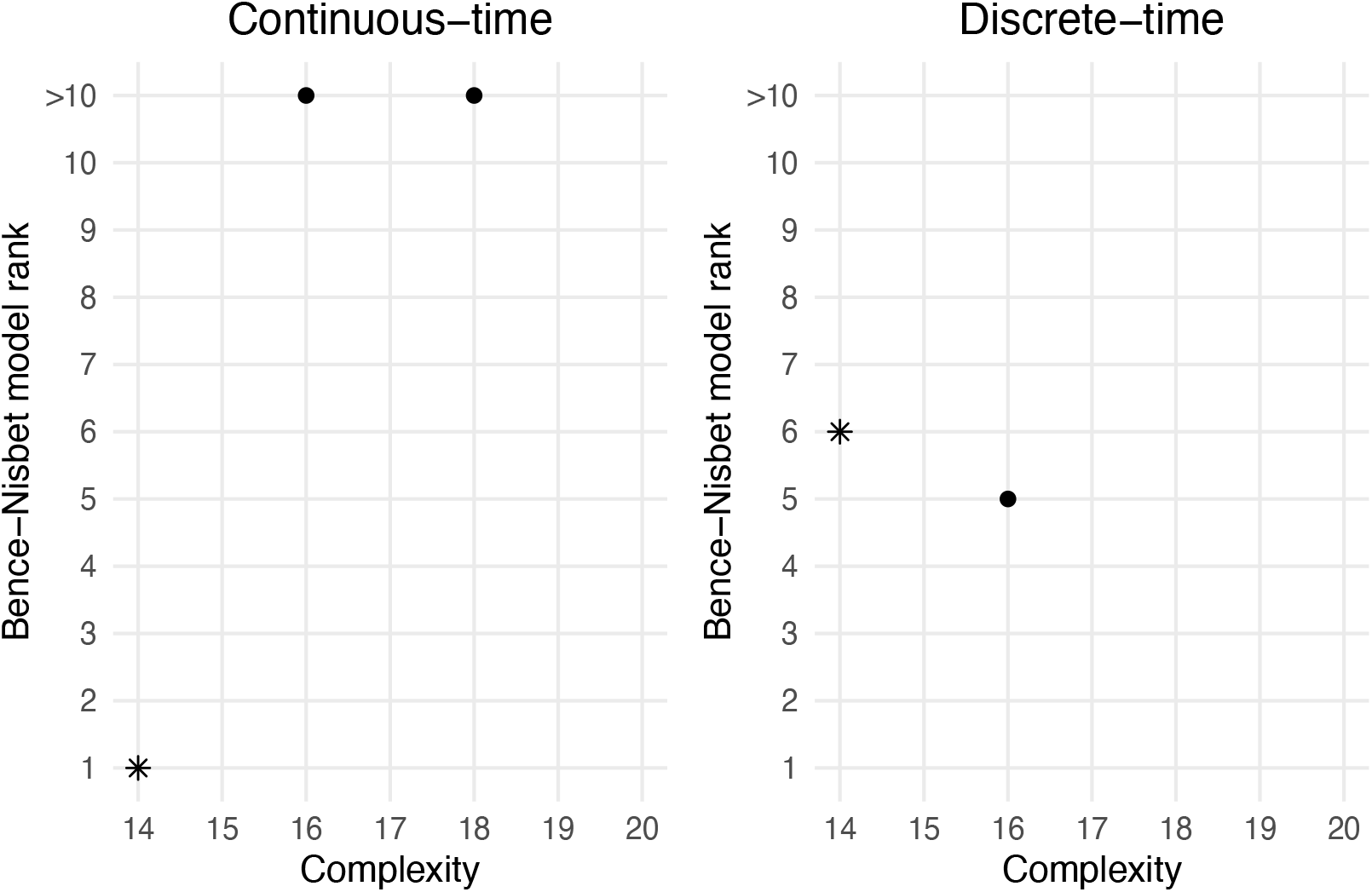
The maximum MSE-rank of the Bence-Nisbet model when it was recovered, returned by the 100 runs from the asymmetric, 100 points per cycle data, with a continuous-time and discrete-time approach, respectively. We assessed the complexity range from 10 to 20. The lowest complexity that the Bence-Nisbet model occurred was complexity 14, which is the complexity level of the Bence-Nisbet model is 14 (indicated by asterisks). Ranks at complexities greater than 14 (indicated by points) occurred when the equations could be algebraically simplified to the Bence-Nisbet model.

### F Attractor reconstruction

**Figure S.3:**
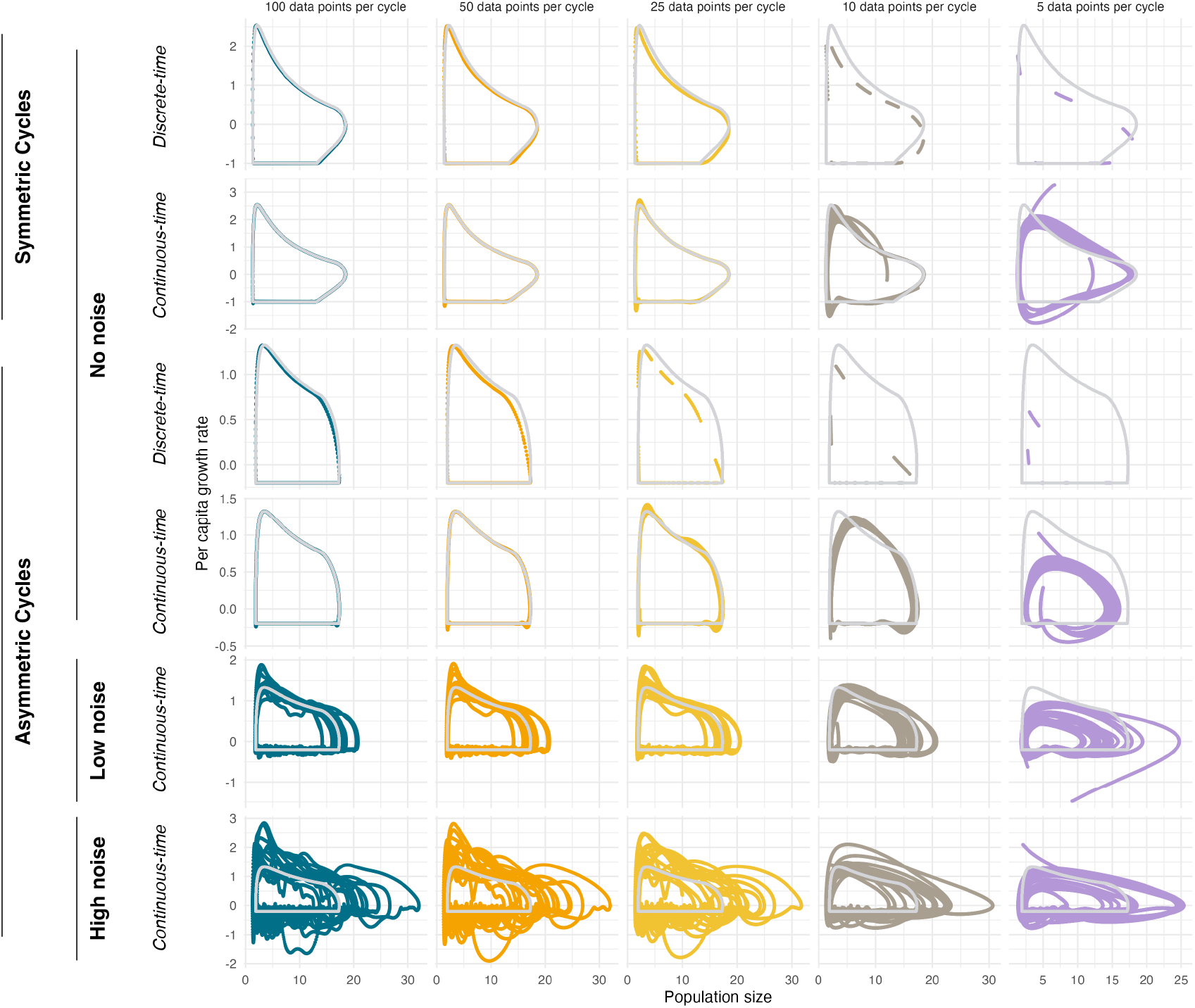
Phase portraits of population size and per capita growth rate (log ratio or interpolated for discrete-time or continuous-time, respectively) provide insight into the influence of sampling density on symbolic regression”s success. As sampling density decreases, the apparent attractor tends to shrink relative to the true attractor of the Bence-Nisbet model (overlaid in light gray), except for the cases with process noise.

